# Biological activities and metabolic profile of endophytic *Bacillus haynesii* from *Momordica charantia*

**DOI:** 10.1101/2024.05.16.594475

**Authors:** Garima Sharma, Surbhi Agarwal, Rashmi Bhardwaj, Vitthal T. Barvkar, Vinay Kumar, Vartika Mathur

**Author notes:** **Correspondence**: Vartika Mathur, Department of Zoology, Sri Venkateswara College, Benito Juarez Marg, Dhaula Kuan, New Delhi-110021.

## Abstract

Endophytes share a complex intimate relationship due to which they are capable of synthesizing metabolites that are either similar or derivative of host bioactive compounds. Such endophytes especially from medicinal plants can be prospected for commercial-scale production of the therapeutic compound. Thus, the study evaluates the antioxidant and anti-inflammatory potential of *Momordica charantia* fruit endophyte *Bacillus haynesii* R2MCFF61, and determines its metabolite profile through LC-Q-TOF-MS for the bioactive compounds produced. *B. haynesii* R2MCFF61 showed significant antioxidant and anti-inflammatory potential with increased DPPH scavenging activity in the standard compound and inhibition of protein denaturation respectively. The metabolic profile of this endophyte showed 23 compounds, mostly consisting of metabolites from lipid, phenylpropanoid and triterpenoid pathways. *B. haynesii* R2MCFF61 showed production of cucurbitane-triterpenoids namely Momordicoside K, 19-dimethoxycucurbita-5(10),6,22(E),24-tetraen-3β-ol and 23(E)-7β-methoxycucurbita-5,23,25-trien-3β-ol. These compounds are mainly produced by *M. charantia* fruit and their production by its endophyte is reported for the first time. Thus, these results indicate the promising therapeutic potential of *B. haynesii* R2MCFF61 which can be utilized as a low-cost sustainable source of these bioactive compounds.

## 1. INTRODUCTION

Cellular homeostasis is disrupted by certain physiological and pathological conditions, increasing reactive oxygen species (ROS), including O_2_^-^ and OH^-^[1]. Oxidative stress conditions are closely linked with the progression of several ailments such as cancer, cardiovascular disease, rheumatoid arthritis, arteriosclerosis, Alzheimer’s and other neurodegenerative disease [2]. Furthermore, ROS can play a regulatory role in activating pro-inflammatory cytokines and inflammasomes while conversely, excessive activation of the inflammatory response can stimulate ROS synthesis [2, 3]. Since natural bioactive metabolites are considered safer and more suitable options, there is an unprecedented increase in the commercial demand for such compounds [4]. Consequently, it becomes necessary to seek sustainable sources of potent and effective bioactive compounds that can provide low-cost therapeutic options.

Endophytes are mutualistic microorganisms that reside inside the plant tissues without causing pathogenesis. These microbes share a complex intimate relationship where the endophytes produce diverse chemicals that promote plant growth and defence [5, 6]. In recent years endophytes, especially from medicinal plants, have received increased attention due to their ability to produce a plethora of natural compounds including steroids, phenols, alkaloids, isocoumarins and terpenoids [7–9]. The main reason for the paradigm shift from medicinal plants to alternate sources is that the growth and output of the plants are more prone to environmental constraints [10]. Moreover, many phytochemicals are produced in low concentrations (depending on plant age and physiology) and have to be extracted from a complex mixture of stereochemicals via specialised methods [11]. In contrast, when their endophytes are provided with suitable conditions such as appropriate media, optimum nutrient availability/limitation and conducive growth parameters, they have the potential to become unlimited resources for a particular compound of pharmaceutical importance [12]. Interestingly, due to the adaptation in the corresponding host, endophytes demonstrate the capacity to synthesize metabolites that closely resemble those found within the host [13, 14]. Thus, such endophytes from medicinal plants can be prospected for commercial-scale production of the therapeutic compound(s).

*Momordica charantia* L., commonly known as the bitter gourd is a member of the Cucurbitaceae family and is known for its bitter-flavoured fruit [15]. It is extensively cultivated in tropical regions for its dual benefits in nutrition and health, as recognized in both Chinese medicinal and Ayurveda approaches. So far, over 228 bioactive compounds, with therapeutic benefits in ailments such as inflammation, ulcers, cancer and diabetes, have been identified from this plant [16, 17]. Additionally, studies have extensively documented the diversity of endophytes inhabiting *M. charantia* and their efficacy in plant stress management [18–21]. However, only a few reports suggest the pharmaceutical potential with anti-diabetic properties by its endophytes [22, 23].

Therefore, the current study aims to evaluate the antioxidant and anti-inflammatory potential of *M. charantia* endophyte *B. haynesii* R2MCFF61. The metabolite profile of the endophyte isolate was evaluated through LC-Q-TOF-MS analysis to determine the bioactive compounds produced. The results of this study are not only important from the pharmaceutical point of view but also provide insight into host-endophyte metabolic interaction.

## 2. MATERIAL AND METHOD

### 2.1. Isolation and characterization of endophyte

*M. charantia* var. Pusa Aushadhi seeds were procured from the Division of Vegetable Science at the Indian Agricultural Research Institute in New Delhi, India. The seeds were sown in garden soil at Sri Venkateswara College (28.5894° N and 77.1681° E), New Delhi and were regularly watered every fourth day along with 1P Hoagland solution every 10 days. Endophyte isolation was done from 50-day old plants, using five fruits per plant in three replicates. The samples were immediately transferred to ice in plastic zip-lock bags and the isolation was done within an hour of sample collection. The fruits were pre-washed with double distilled water (DDW) to remove dust particles. The isolation of endophyte was done as per Sharma et al. [24] using one gram of the surface sterilized fruit sample and 10^-6^ times diluted extract was inoculated on Reasoner’s 2 agar (R2A; Himedia Laboratories, India). The R2A plates were incubated at 30 °C for seven days and were observed every 24h. Based on the morphology three microbial colonies were obtained and respective colony-forming units were counted.

For identification, the pure culture of the isolate was obtained by two rounds of colony sub-culture. This pure culture was then inoculated in 10ml of nutrient broth and allowed to incubate overnight at a temperature of 30°C. The inoculum was centrifuged at 8500rpm at 4°C for 20 min. The supernatant was discarded and the subsequent pellet was used for DNA extraction through the CTAB method [25]. The DNA sample was used for Sanger sequencing using the universal primers targeting the 16S rRNA region. By aligning the forward and reverse sequences thus generated a consensus sequence was created using Finch TV version 1.4.0 and BioEdit version 7.1.3.0 software via the “cap contig assembly program”. The Consensus sequence was compared to the NCBI database and a phylogenetic analysis with 25 closely related sequences was then constructed using MEGA (Version 11) software, employing the neighbour-joining method with bootstrap analysis (1000 repeats) [26].

### 2.2. Preparation of cell-free extract

Pure culture of the isolate was cultured in nutrient broth (Himedia Laboratories, India) for a duration of 21 days at 30°C and 150 rpm as per Shweta et al. [27]. To separate the cells, the culture was centrifuged at 8000rpm at 4°C for 20 min. The supernatant was collected and designated as Aqueous extract (AqE), whereas the pellet was re-suspended in 60% ethanol for overnight at RT. Subsequently, the pellets were subjected to eight cycles of sonication at 70% output amplitude of 130W and 20kHz for 20 seconds pulse and a rest period of 1 min on ice. To remove the cell debris, the pellet extract was centrifuged at 10,000 rpm for 15 min, and the supernatant was collected and marked as cell-free extract (CFE). Both CFE and AqE were filtered via a 0.2 µm filter (Millipore ®, Merck) and lyophilized for further analysis.

### 2.3 Estimation of total phenol

From the lyophilized extracts, two mg/ml stock was prepared in 60% ethanol for total phenol estimation using the method by Makkar et al. [28]. For total phenols, 100μl extract was mixed with 1.25ml of 20% sodium carbonate and 250μl 1N Folin-Ciocalteu reagent (SRL Pvt. Ltd., India). This reaction mixture was incubated at RT in the dark for 45 min and the blue-colored complex was measured at 725nm using a UV Spectrophotometer (UV1800, Shimadzu, Japan). Blank consisted equal quantity of 60% ethanol instead of the stock solution. The total phenol content (mg/g extract) was determined by comparing it with a standard solution of tannic acid (0.1mg/mL, SRL Pvt. Ltd., India).

### 2.4 Antioxidant activity

The antioxidant potential of CFE and AqE was assessed by the 2,2-diphenyl-2-picrylhydrazyl (DPPH) decolorization method described by Velho-Pereira et al. [29]. Using the lyophilized extract 10mg/ml stock was prepared in 60% ethanol. Three-millilitre reaction mixture constituted of 20 μl CFE or 200 μl AqE in 1.5ml 0.1mM DPPH and the rest of the volume was made up by absolute ethanol. Potential antioxidant activity was determined by the reduction in absorbance at 517nm. The ability to scavenge free radicals was calculated using the following formula:

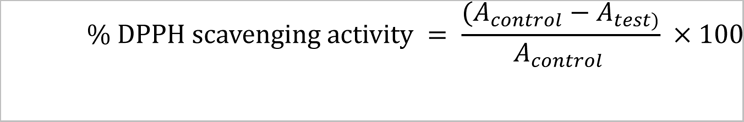

### 2.5 Anti-inflammatory activity

The anti-inflammatory activity was evaluated by the inhibition of protein denaturation method described by Senthilkumar et al. [30]. The test reaction (T_extract_) consisted of 0.45ml 5% w/v bovine serum albumin (BSA) solution and 0.05ml of the test extract (10mg/ml CFE or AqE stock), while the control reaction (T_control_) contained 0.05 ml DDW (instead of extract). Using different concentrations (50, 100, 150, 200, 250 μg/ml) of diclofenac sodium, a standard curve was prepared in 0.45 ml of the 5% w/v BSA solution. The absorbance was measured at 416nm and the percentage inhibition was calculated using the following equation, where T_control_ represented 100% protein denaturation:

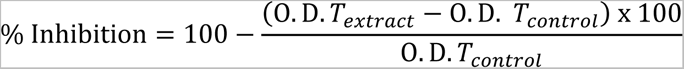

### 2.6 Metabolite profiling using liquid chromatography quadrupole time of flight mass spectrometry

The sample extraction and metabolic profile through liquid chromatography quadrupole time-of-flight mass spectrometry (LC-Q-TOF-MS) (in both positive and negative modes) was performed as per Vasav et al. [31]. 100mg lyophilized of AqE was dissolved in 100% MS grade Methanol and vortexed for 10min at RT. The sample was then sonicated for 20 min, centrifuged at 13000rpm for 10min and filtered with a 0.22 µm nylon syringe filter. The sample was loaded on the Infinity II 1260 LC system equipped with an Infinity Lab Poroshell 120 EC-C18 column (2.1 × 150 mm, 1.9 μm particle size, Agilent, USA) maintained at 40 °C, for metabolite separation. The mobile phase consisted of 100% MS grade water (Solvent A) and 100% MS grade acetonitrile (Solvent B), both containing 0.1% formic acid, with a flow rate of 0.3 ml/min and 1.5 μl injection volume. A gradient of the mobile phase was used for 20 minutes with 98% solvent A and 2% Solvent B for 0.3 min in the initial phase. The gradient gradually reached 30% at 2 min, 45% at 7 min, and finally reaching 98% at 12 minutes, followed by a three-minute hold. In the last five minutes of the run, the column was equilibrated with a ratio of 98%:2%. Data was acquired with Agilent 6530 LC-Q-TOF mass spectrometer (Agilent, USA) pre-calibrated with ESI-L low concentration tuning mixture (Part No G1969-85000, Agilent, USA). Deisotoping, deconvolution, peak picking and retention time correction were performed with default parameters [32]. Ion chromatograms from the extended dynamic range data were extracted using MassHunter Workstation software (B.08.00, Agilent). Peak area and fold change was estimated by Agilent mass-hunter qualitative workflow B.08.00.

### 2.7 Statistics

Normality of the data and homogeneity of variance were analysed by Kolmogorov–Smirnov test and Levene’s test respectively using SPSS 23 (IBM Chicago, IL, USA). The data of the antioxidant assay was normalized by log transformation. Antioxidant and anti-inflammatory assays were evaluated using univariate ANOVA with Tukey post-hoc.

## 3. RESULTS AND DISCUSSION

### 3.1. Isolation and characterization of endophyte

The isolate R2MCFF61 showed a white dry texture, circular colonies with light uneven margins and a raised centre on R2A (Fig 1a). It showed a slow growth rate and constituted about 6 (± 2.53) x 10^5^ CFU/ml per gram of fresh fruit. The isolate was identified as *Bacillus haynesii* (GenBank Assession no. PP565064) by 16S rRNA sequencing and exhibited 99% similarity with *B. haynesii* strain NRRL B-41327. The phylogenetic tree depicting relatedness to 20 closest neighbours is given in Fig. 1b.

**Figure 1:**
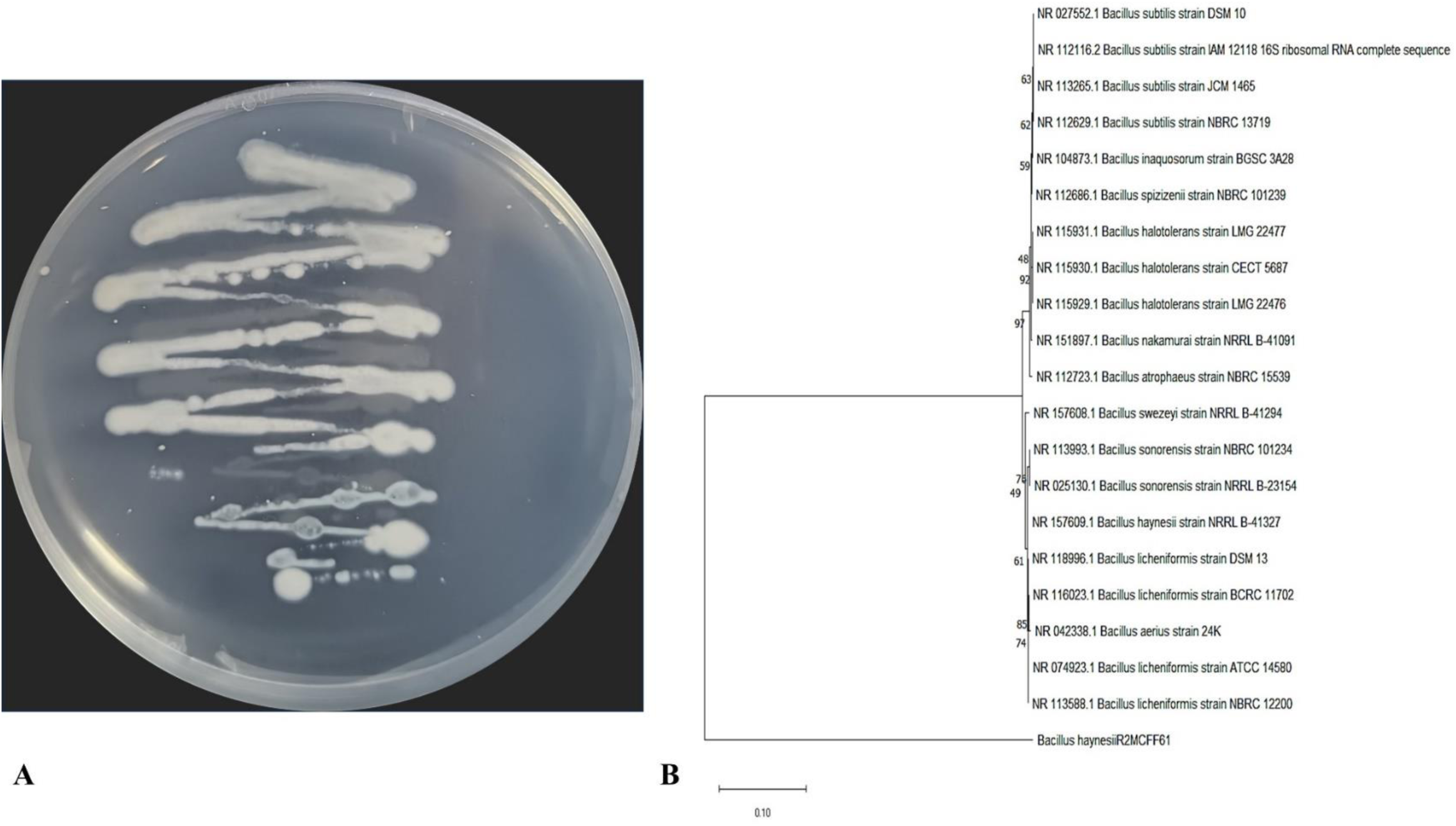
Isolation and Identification of *Momordica charantia* endophyte *Bacillus haynesii* R2MCFF61 (a) Pure cultures of the endophytic isolated; (b) Phylogenetic tree of 16S rRNA sequences, conducted with MEGA 10 using neighbor-joining method with bootstrap value (1000 replicates)

to date, only a limited number of studies have documented bacterial endophytes of *M. charantia* primarily associated with the roots, where the genus *Bacillus* is predominant, constituting *B. subtilis* and *B. licheniformis* [19, 21]. *Bacillus* stands out as one of the most prevalent genera amongst bacterial endophytes, playing dual roles in promoting plant growth and fortifying defences against pathogens [33, 34]. Additionally, strains of *B. haynesii* have been identified as endophytes associated with medicinal plants such as Black cumin and Holy Basil [35, 36], and exhibiting the capability to induce resistance against phytopathogens in their respective hosts [35, 37].

### 3.2. Estimation of total phenol

With 3.35 ± 0.06 mg/g the CFE of *B. haynesii* R2MCFF61 contained higher phenolic compounds compared to its AqE (2.5 ± 0.23 mg/g), indicating their limited secretion into the extracellular matrix. The fruit of *M. charantia* is rich in phenol compounds, reported in the range of 1.5 – 9 mg/gm [38]. Phenol compounds are the secondary metabolites involved in free radical scavenging and thus are responsible for the antioxidant potential of the plant. Phenolic metabolites comprise of a diverse group of compounds, their biosynthesis occurs mainly via phenylpropanoids and the mevalonate pathway [39].

### 3.3. Antioxidant activity

CFE of the endophyte *B. haynesii* R2MCFF61 showed significantly increased DPPH scavenging activity than both the standard compound as well as its AqE indicating better antioxidant potential than the standard compound (univariate ANOVA; *F*_(2,8)_= 23.508, *P*<0.05) (Fig 2).

**Figure 2:**
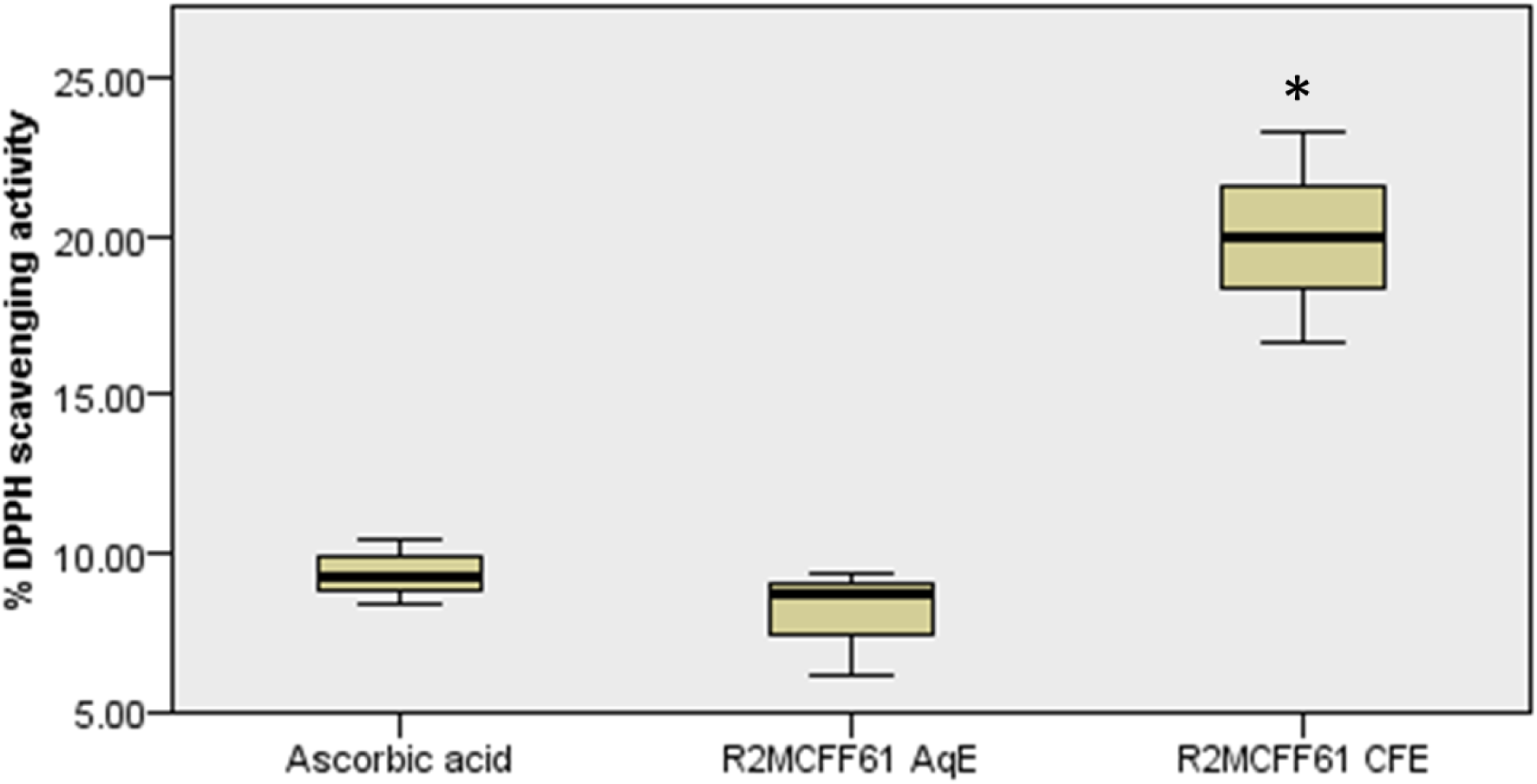
Percentage DPPH scavenging activity (± SE) of *Momordica charantia* endophyte *Bacillus haynesii* R2MCFF61 (n=3)

For establishment and survival within their host, endophytes synthesize antioxidant molecules capable of alleviating the oxidative stress generated inside the host tissues [40]. These metabolic adaptations enable many endophytes to produce a variety of antioxidant compounds tailored to the specific host environment [41, 42]. Total phenol content in endophytes positively correlates with their antioxidant capacity [43], a trend also observed in our study of the CFE from the endophyte. Notably, *B. haynesii* also showed antioxidant potential in association with other medicinal plants such as Black Cumin [35]. Nevertheless, limited information is reported regarding the metabolite profile of this *B*. *haynesii* during endophyte association.

### 3.4. Anti-inflammatory activity

The test extracts were at par with the standard compound and showed a significant difference in the protein denaturation activity (univariate ANOVA; *F*_(2,8)_= 1.469, *P*=0.303). Moreover, no significant difference was observed between the AqE and CFE (Fig 3). Protein denaturation in the tissues is a well-known cause of inflammation. Consequently, compounds such as salicylic acid and diclofenac sodium, which aids in preventing this process, are employed as anti-inflammatory drugs [44, 45]. Extracts from endophytes such as *Aspergillus* sp., *Penicillium* sp., *Rhizopus* and *Bacillus subtilis* show anti-inflammatory potential which ranged between 30-87% [44, 46]. *B. haynesii* R2MCFF61 extract showed more than 100% denaturation capacity at the concentration of 50 μg/ml.

**Figure 3:**
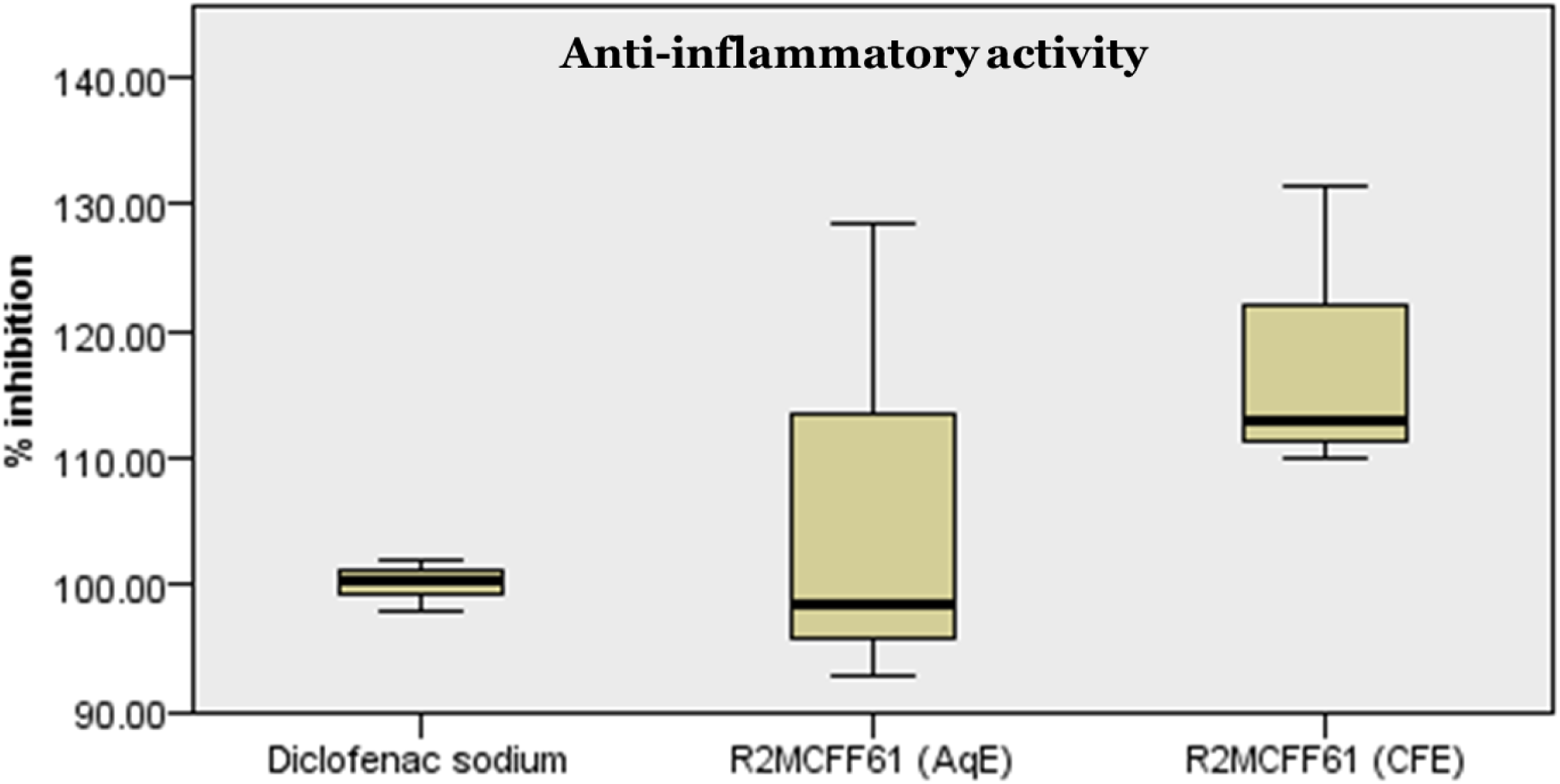
Percentage inhibition (± SE) of protein denaturation (anti-inflammatory) activity of *Momordica charantia* endophyte *Bacillus haynesii* R2MCFF61 (n=3)

### 3.5. Metabolite profiling using liquid chromatography quadrupole time of flight mass spectrometry

Endophytes residing in medicinal plants thrive in a highly competitive environment by adapting to bioactive metabolites. Through the intricate interactions, they produce analogous or derivatives of the host-specific molecules even in isolated culture systems [47]. Endophytes, particularly those belonging to the genus *Bacillus*, have been reported to synthesize a diverse range of secondary metabolites such as iturin, surfactin, bacillibactin and bacillomycin [48, 49]. However, *B. haynesii* has garnered significant attention for its enzymatic capabilities, especially in the production of enzymes such as L-methioninase, acid phosphatase and Keratinase, with limited reports focusing on its metabolic profile [50–53].

In our study, the fruit endophyte *B. haynesii* R2MCFF61 CFE showed the presence of 23 compounds, mostly consisting of lipid, phenylpropanoid and triterpenoid metabolites, out of these 12 are previously reported for bioactive properties. The most abundant compounds of the extract in terms of peak area intensity were 24-nor-5beta-cholane-3alpha,7alpha,12alpha,23-tetrol (2.82 x 10^7^), cycloleucine (2.02 x 10^7^) and hexadecanoic acid (2.02 x 10^7^) (detailed profile of the metabolites is provided in Table 1). However, this is the first report of *B. haynesii* strain to produce triterpenoid compounds such as Momordicoside K, 19-dimethoxycucurbita-5(10),6,22(E),24-tetraen-3β-ol and 23(E)-7β-methoxycucurbita-5,23,25-trien-3β-ol. These three compounds are specific to the family Cucurbitaceae and have been previously reported in *M. charantia* fruit. Momordicoside K is a known anticancer agent, whereas the other two compounds show antioxidant activity [54, 55]. Chromatograms of the peaks of bioactive metabolites are provided in Fig 4 a & b.

**Table 1:** Metabolite profiling of the *Momordica charantia* endophytes *Fusarium* sp. P2MCLC3*, Alcaligens faecalis* R2MCL41 and *Escherichia fergusonii R2MCFF51* using LC-Q-TOF-MS.

| S. No. | Name | Formula | m/z | Mass | Diff<br>(Tgt,<br>ppm) | Score | RT | Area | Ions | Activity |
| --- | --- | --- | --- | --- | --- | --- | --- | --- | --- | --- |
| 1 | Benzoic acid | C<sub>7</sub>H<sub>6</sub>O<sub>2</sub> | 140.07 | 122.04 | 1.21 | 87.37 | 1.25 | 2.24E+05 | 2 | Antimicrobial |
| 2 | 1,3,7-Trimethyluric acid | C<sub>8</sub>H<sub>10</sub>N<sub>4</sub>O<sub>3</sub> | 233.07 | 210.07 | -7.11 | 62.94 | 1.27 | 1.91E+05 | 5 | Antioxidant |
| 3 | 5-Aminovaleric acid | C<sub>5</sub>H<sub>11</sub>NO<sub>2</sub> | 140.07 | 117.08 | 3.91 | 57.79 | 1.46 | 1.57E+07 | 3 | - |
| 4 | Cycloleucine | C<sub>6</sub>H<sub>11</sub>NO<sub>2</sub> | 147.11 | 129.08 | 4.69 | 84.75 | 1.6 | 2.02E+07 | 2 | Immunosuppressive;<br>Antineoplastic;<br>Cytostatic |
| 5 | Neuraminic acid | C<sub>9</sub>H<sub>17</sub>NO<sub>8</sub> | 268.10 | 267.10 | -0.34 | 74.96 | 1.79 | 1.02E+07 | 2 | - |
| 6 | 2-Piperidone<br>(aka δ-Valerolactam) | C<sub>5</sub>H<sub>9</sub>NO | 100.08 | 99.07 | 6.72 | 89.15 | 3.62 | 3.79E+06 | 3 | Anticancer;<br>Antioxidant; Anti-inflammatory |
| 7 | Bluemenol A | C<sub>13</sub>H<sub>20</sub>O<sub>3</sub> | 263.10 | 224.14 | 1.71 | 63.46 | 4.16 | 2.12E+06 | 2 | Anticancer |
| 8 | Ethyl-4,5-dimethylphenol | C<sub>14</sub>H<sub>22</sub>O | 245.13 | 206.17 | 1.66 | 74.05 | 4.18 | 1.08E+07 | 3 | - |
| 9 | 19-dimethoxycucurbita-5(10),6,22(E),24-tetraen-3β-ol | C<sub>32</sub>H<sub>50</sub>O<sub>3</sub> | 505.36 | 482.37 | -4.84 | 53.17 | 9.81 | 3.74E+06 | 2 | Antioxidant |
| 10 | Hexadecanoic acid | C<sub>16</sub>H<sub>32</sub>O<sub>2</sub> | 257.25 | 256.24 | 5.74 | 82.95 | 10.41 | 2.02E+07 | 8 | - |
| 11 | Xestoaminol C | C<sub>14</sub>H<sub>31</sub>N O | 230.25 | 229.24 | 3.45 | 63.82 | 10.52 | 7.10E+05 | 4 | Anticancer |
| 12 | Sphinganine | C<sub>18</sub>H<sub>39</sub>NO<sub>2</sub> | 302.31 | 301.30 | 6.42 | 71.82 | 12.11 | 9.65E+05 | 8 | - |
| 13 | Linoleamide | C<sub>18</sub>H<sub>33</sub>NO | 302.25 | 279.26 | 7.19 | 74.63 | 13.62 | 2.09E+05 | 4 | - |
| 14 | Proanthocyanidin A2 | C<sub>30</sub>H<sub>24</sub>O<sub>12</sub> | 594.16 | 576.13 | 5.76 | 76.07 | 14.03 | 4.46E+04 | 10 | Inflammatory;<br>Antioxidant;<br>Antiviral |
| 15 | Momordicoside K | C<sub>37</sub>H<sub>60</sub>O<sub>9</sub> | 666.47 | 648.42 | -2.87 | 59.13 | 15.01 | 6.22E+06 | 5 | Anticancer |
| 16 | Linoleic acid | C<sub>18</sub>H<sub>32</sub>O<sub>2</sub> | 298.28 | 280.24 | -1.11 | 52.73 | 15.37 | 1.15E+07 | 6 | - |
| 17 | Oleic acid | C<sub>18</sub>H<sub>34</sub>O<sub>2</sub> | 283.26 | 282.25 | -3.39 | 78.42 | 16.59 | 1.01E+07 | 8 | - |
| 18 | Cis-9-hexadecenal | C<sub>16</sub>H<sub>30</sub>O | 239.24 | 238.23 | -5.9 | 60.04 | 17.16 | 1.08E+06 | 2 | Antifungal |
| 19 | 11-amino-undecanoic acid | C<sub>11</sub>H<sub>23</sub>NO<sub>2</sub> | 219.21 | 201.17 | 1.61 | 62.97 | 17.17 | 2.00E+05 | 7 | - |
| 20 | Elaidamide | C<sub>18</sub>H<sub>35</sub>NO | 282.28 | 281.27 | 4.17 | 56.73 | 17.36 | 8.27E+05 | 9 | - |
| 21 | (23E)-7β-methoxycucurbita-5,23,25-trien-3β-ol | C<sub>31</sub>H<sub>50</sub>O<sub>2</sub> | 477.37 | 454.38 | -6.65 | 56.99 | 18.05 | 6.20E+05 | 2 | Antioxidant |
| 22 | 25,26,27-trinorcucurbit-5-ene3,7,23-trione | C<sub>27</sub>H<sub>40</sub>O<sub>3</sub> | 430.33 | 412.30 | 7.35 | 56.33 | 18.08 | 4.15E+06 | 3 | Antioxidant |
| 23 | 24-Nor-5β-cholane-3α,7α,12α,23-tetrol | C<sub>23</sub>H<sub>40</sub>O<sub>4</sub> | 381.30 | 380.29 | 1.1 | 70.09 | 18.63 | 2.82E+07 | 3 | - |

**Figure 4a:**
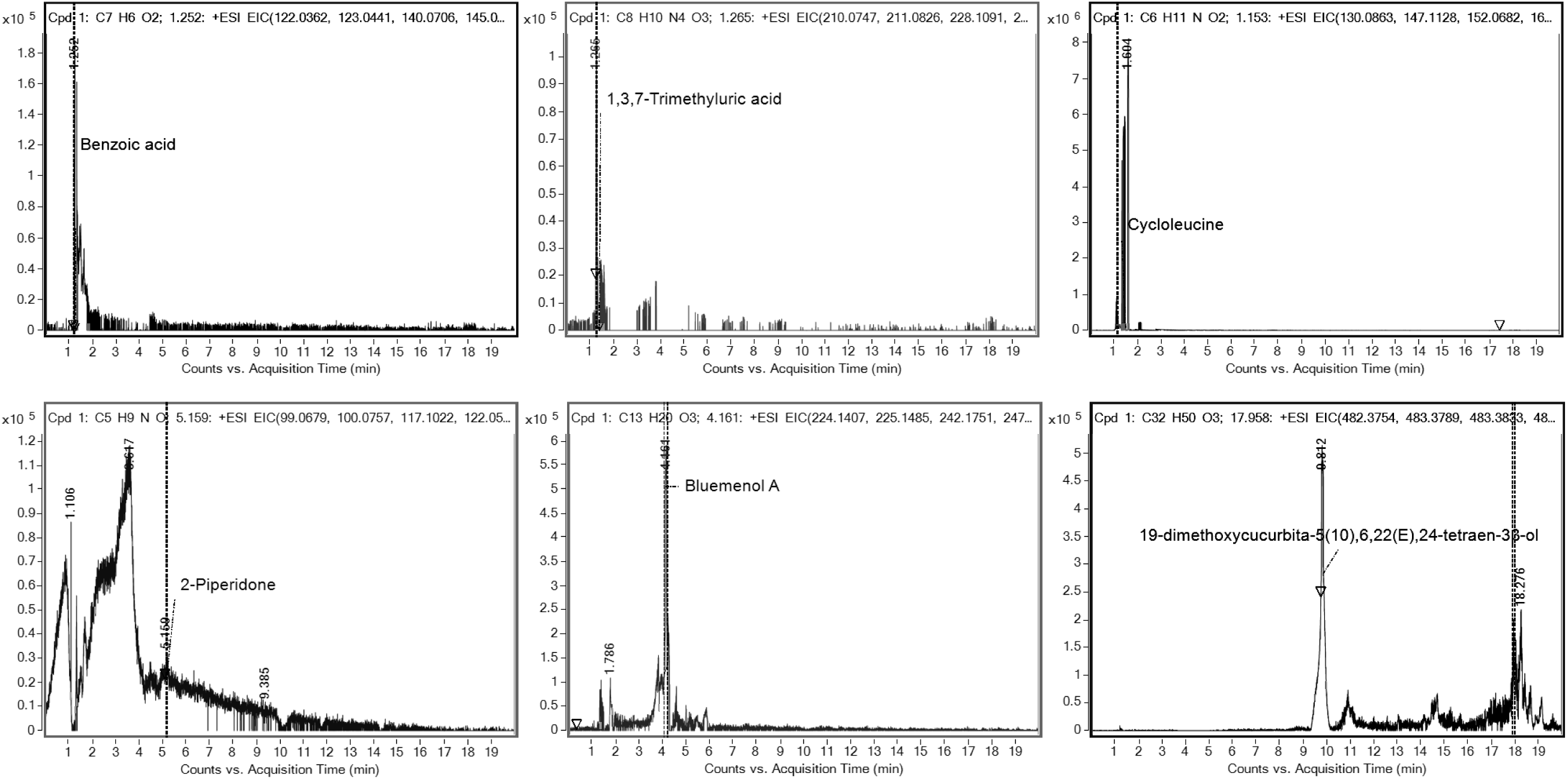
Representative LC-MS chromatographic peaks of the compounds Benzoic acid, 1,3,7-Trimethyluric acid, Cycloleucine, 2-Piperidone, Bluemenol A and 19-dimethoxycucurbita5(10),6,22(E),24-tetraen-3β-ol from *Momordica charantia* endophytes *Bacillus haynesii* R2MCFF61. Retention tim444e in minutes is written on the top of the respective peak

**Figure 4b:**
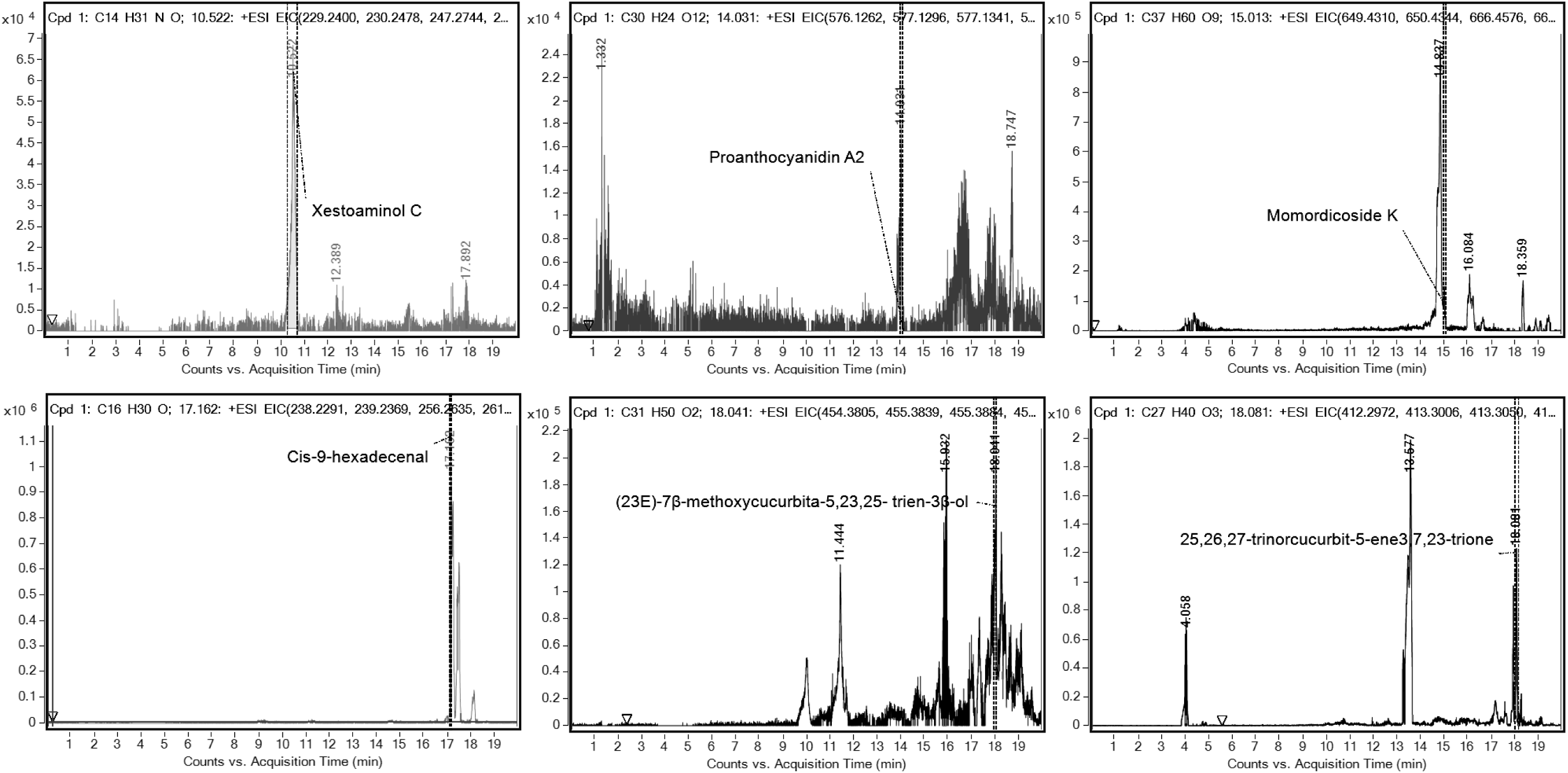
Representative LC-MS chromatographic peaks of the compounds Xestoaminol C, Proanthocyanidin A2, Momordicoside K, Cis-9-hexadecenal, (23E)-7β-methoxycucurbita-5,23,25-trien-3β-ol and 25,26,27-trinorcucurbit-5-ene3,7,23-trione from *Momordica charantia* endophytes *Bacillus haynesii* R2MCFF61. Retention time in minutes is written on the top of the respective peak

The compound observed with the second highest peak area intensity, Cycloleucine (also known as 1-aminocyclopentane carboxylic acid) is also reported from *M. charantia* fruit and is known for its anti-inflammatory, cytotoxic, immunosuppressive and antimicrobial properties [56, 57]. Besides these, antioxidant compounds such as 1,3,7-trimethyluric acid, 2-piperidone and proanthocyanidin A2 were also observed in this extract. While 1,3,7-trimethyluric acid mainly acts as a mild antioxidant, Proanthocyanidin A2 is reported for antioxidant, anti-inflammatory and antiviral properties [58, 59]. However, the smallest peak area intensity of proanthocyanidin A2 indicates its limited production by *B. haynesii* R2MCFF61. 2-piperidone is a well-known bioactive compound with antioxidant, anti-inflammatory and anticancer properties [60]. Although this compound is mainly produced in the host fruit, it has also been reported from a different endophytic *Bacillus* sp. [61]. This extract also showed the presence of two anticancer compounds namely Bluemenol A and xestoaminol C [62, 63].

Antimicrobial compounds such as benzoic acid and cis-9-hexadecenal were also observed in the extract. While benzoic acid itself shows limited toxicity, its derivatives are important antimicrobial agents [64, 65]. In contrast, cis-9-hexadecenal is reported for antifungal activity against *Aspergillus fumigatus* [66]. Several studies report the antimicrobial activity of *B. haynesii* strains isolated from variety of sources such as plants and marine soil [35, 67]. These results indicate a possibility of utilizing the endophyte for the production of such antimicrobial compounds. The presence of such compounds gives a competitive advantage to the endophyte for survival and defence inside the host plant [47]. The extract also consisted of lipid compounds such as linoleic, oleic and hexadecanoic acid (with high peak area intensities) as well as a fatty acid amide derivative namely linoleamide. The pattern of lipid components in *Bacillus* is essential for survival and adaptation to a range of environmental conditions [68]. Furthermore, amide derivatives of fatty acids are essential for self-defence in endophytes [69]. Another, compound with high peak area intensity observed in this endophyte extract of is neuraminic acid (or sialic acid). Sialic acid and its derivatives are important for the survival of the microbe inside the host organism through functions such as receptor adherence and biofilm production [70]. Thus, the specific pattern of these compounds is possibly essential for the adaptation of this endophyte to its host.

## 4 CONCLUSION

*B. haynesii* R2MCFF61 is one of the predominant culturable slow growing endophytes isolated from the fruit of *M. charantia*. The extract derived from this endophyte showed significant antioxidant and anti-inflammatory properties which may be attribute to the presence of twelve bioactive compounds, primarily comprising terpenoid and phenylpropanoid metabolites. Notably, the endophyte demonstrated a remarkable synchrony in metabolite production with those typically found in the bitter gourd fruit. Accordingly, this is the first report of endophyte that produce cucurbitane-triterpenoids momordicoside K, 19-dimethoxycucurbita-5(10),6,22(E),24-tetraen-3β-ol and 23(E)-7β-methoxycucurbita-5,23,25-trien-3β-ol which is originally reported in *M. charantia* fruit. Future studies may focus on the upscale production and extraction of these compounds, potentially utilizing this endophyte as a promising low-cost sustainable source of these bioactive compounds.

## Acknowledgement

The authors would like to thank the research trainee Kanishka Kumar (Animal Plant Interactions Lab) for her help during various parts of the experiments. The authors would also like to thank Dr. Ravindra Varma Polisetty, Department of Biochemistry, Sri Venkateswara College, University of Delhi, India, for providing access to their sonicator system.

## Reference

1. Valko M, Leibfritz D, Moncol J, Cronin MT, Mazur M, Telser J. Free radicals and antioxidants in normal physiological functions and human disease. Int J Biochem Cell Biol. 2007; 39(1):44–84. Epub 2006/09/19. doi: 10.1016/j.biocel.2006.07.001.

2. Zuo L, Prather ER, Stetskiv M, Garrison DE, Meade JR, Peace TI, Zhou T. Inflammaging and oxidative stress in human diseases: from molecular mechanisms to novel treatments. Int J Mol Sci. 2019; 20(18):4472.

3. Biswas SK. Does the Interdependence between Oxidative Stress and Inflammation Explain the Antioxidant Paradox? Oxid Med Cell Longev. 2016; 2016:5698931. doi: 10.1155/2016/5698931. PubMed PMID: 26881031

4. Braga A, Rocha I, Faria N. Microbial Hosts as a Promising Platform for Polyphenol Production. In: Akhtar MS, Swamy MK, Sinniah UR, editors. Natural Bio-active Compounds: Volume 1: Production and Applications. Singapore: Springer Singapore; 2019. p. 71–103.

5. Aly A, Debbab A, Proksch P. Fungal endophytes–secret producers of bioactive plant metabolites. Pharmazie. 2013; 68(7):499–505.

6. Wani ZA, Ashraf N, Mohiuddin T, Riyaz-Ul-Hassan S. Plant-endophyte symbiosis, an ecological perspective. Appl Microbiol Biotechnol. 2015; 99(7):2955–65.

7. Knight V, Sanglier JJ, DiTullio D, Braccili S, Bonner P, Waters J, Hughes D, Zhang L. Diversifying microbial natural products for drug discovery. Appl Microbiol Biotechnol. 2003; 62(5):446–58. doi: 10.1007/s00253-003-1381-9.

8. Kusari S, Hertweck C, Spiteller M. Chemical ecology of endophytic fungi: origins of secondary metabolites. Chem Biol. 2012; 19(7):792–8.

9. Hashem AH, Al-Askar AA, Abd Elgawad H, Abdelaziz AM. Bacterial Endophytes from Moringa oleifera Leaves as a Promising Source for Bioactive Compounds. Separations. 2023; 10(7).

10. Ogbe AA, Finnie JF, Van Staden J. The role of endophytes in secondary metabolites accumulation in medicinal plants under abiotic stress. S Afr J Bot. 2020; 134:126–34. doi: 10.1016/j.sajb.2020.06.023.

11. Pyne ME, Narcross L, Martin VJ. Engineering plant secondary metabolism in microbial systems. Plant Physiol. 2019; 179(3):844–61.

12. Chathurdevi G, Gowrie S. Endophytic fungi isolated from medicinal plant—a source of potential bioactive metabolites. Int J Curr Pharm Res. 2016; 8:50–6.

13. Aly AH, Debbab A, Kjer J, Proksch P. Fungal endophytes from higher plants: a prolific source of phytochemicals and other bioactive natural products. Fungal Divers. 2010; 41:1–16.

14. Dudeja S, Singh, Giri R. Beneficial properties, colonization, establishment and molecular diversity of endophytic bacteria in legumes and non legumes. Afr J Microbiol Res. 2014; 8(15):1562–72. doi: 10.5897/AJMR2013.6541.

15. Grover JK, Yadav SP. Pharmacological actions and potential uses of Momordica charantia: a review. J Ethnopharmacol. 2004; 93(1):123–32. doi: 10.1016/j.jep.2004.03.035.

16. Gayathry K, John JA. A comprehensive review on bitter gourd (Momordica charantia L.) as a gold mine of functional bioactive components for therapeutic foods. Food Production, Processing and Nutrition. 2022; 4(1):1–14. doi: 10.1186/s43014-022-00089-x.

17. Villarreal-La Torre VE, Guarniz WS, Silva-Correa C, Cruzado-Razco L, Siche R. Antimicrobial activity and chemical composition of Momordica Charantia: A review. Pharmacogn J. 2020; 12(1).

18. Huang J-H, Xiang M-M, Jiang Z-D. Endophytic fungi of bitter melon (Momordica Charantia) in Guangdong province, China. Great Lakes Entomol. 2012; 45(1 & 2):2.

19. Singh R, Pandey KD, Singh M, Singh SK, Hashem A, Al-Arjani A-BF, Abd Allah EF, Singh PK, Kumar A. Isolation and Characterization of Endophytes Bacterial Strains of Momordica charantia L. and Their Possible Approach in Stress Management. Microorganisms 2022; 10(2):290. doi: 10.3390/microorganisms10020290.

20. Etim D, Okon N. Molecular characterization of virus infecting Momordica charantia Linn and the application of Trichoderma viride as Biocontrol Agent in Baccocco Cross River State, Nigeria. Annu Res Rev Biol. 2021;36(12):51–8.

21. Singh R, Kumar A, Singh M, Pandey KD. Isolation and Characterization of Plant Growth Promoting Rhizobacteria From Momordica Charantia L. In: Singh AK, Kumar A, Singh PK, editors. PGPR Amelioration in Sustainable Agriculture: Woodhead Publishing; 2019. p. 217–38.

22. Pavithra N, Sathish L, Babu N, Venkatarathanamma V, Pushpalatha H, Reddy GB, Ananda K. Evaluation of α-amylase, α-glucosidase and aldose reductase inhibitors in ethyl acetate extracts of endophytic fungi isolated from anti-diabetic medicinal plants. Int J Pharm Sci Res. 2014; 5(12):5334–41. doi: 10.13040/IJPSR.0975-8232.

23. Deshmukh SR, Dhas YK, Patil B. Comparative account on medicinal importance of Momordica charantia and its endophytes. World J Pharm Res. 2014; 3:632–40.

24. Sharma G, Rahul, Guleria R, Mathur V. Differences in plant metabolites and microbes associated with Azadirachta indica with variation in air pollution. Environ Pollut. 2020; 257:113595. doi: 10.1016/j.envpol.2019.113595.

25. Wilson K. Preparation of genomic DNA from bacteria. Curr Protoc Mol Biol. 2001; 56(1): 2–4. doi: 10.1002/0471142727.mb0204s56.

26. Tamura K, Stecher G, Kumar S. MEGA11: molecular evolutionary genetics analysis version 11. Mol Biol Evol. 2021; 38(7):3022–7.

27. Shweta S, Bindu JH, Raghu J, Suma H, Manjunatha B, Kumara PM, Ravikanth G, Nataraja KN, Ganeshaiah KN, Uma Shaanker R. Isolation of endophytic bacteria producing the anti-cancer alkaloid camptothecine from Miquelia dentata Bedd.(Icacinaceae). Phytomed. 2013; 20(10):913–7.

28. Makkar HP, Siddhuraju P, Becker K. Plant secondary metabolites. Totowa, NJ, USA: Humana Press; 2007.

29. Velho-Pereira S, Parvatkar P, Furtado IJ. Evaluation of antioxidant producing potential of halophilic bacterial bionts from marine invertebrates. Indian J Pharm Sci. 2015; 77(2):183.

30. Senthilkumar N, Nandhakumar E, Priya P, Soni D, Vimalan M, Potheher IV. Synthesis of ZnO nanoparticles using leaf extract of Tectona grandis (L.) and their anti-bacterial, anti-arthritic, anti-oxidant and in vitro cytotoxicity activities. New J Chem. 2017; 41(18):10347–56.

31. Vasav AP, Pable AA, Barvkar VT. Differential transcriptome and metabolome analysis of Plumbago zeylanica L. reveal putative genes involved in plumbagin biosynthesis. Fitoterapia. 2020;147:104761. doi: 10.1016/j.fitote.2020.104761.

32. Tautenhahn R, Patti GJ, Rinehart D, Siuzdak G. XCMS Online: a web-based platform to process untargeted metabolomic data. Anal Chem. 2012;84(11):5035–9. doi: 10.1021/ac300698c.

33. Afzal I, Shinwari ZK, Sikandar S, Shahzad S. Plant beneficial endophytic bacteria: Mechanisms, diversity, host range and genetic determinants. Microbiol Res. 2019; 221:36–49. doi: 10.1016/j.micres.2019.02.001.

34. Matloub AA, Gomaa EZ, Hassan AA, Elbatanony MM, El-Senousy WM. Comparative chemical and bioactivity studies of intra-and extracellular metabolites of endophytic bacteria, Bacillus subtilis NCIB 3610. Int J Pept Res Ther. 2020; 26(1):497–511.

35. Litao N, Rustamova N, Paerhati P, Ning H-X, Yili A. Culturable Diversity and Biological Properties of Bacterial Endophytes Associated with the Medicinal Plants of Vernonia anthelmintica (L.) Willd. Appl Sci. 2023; 13(17).

36. Sahu PK, Singh S, Singh UB, Chakdar H, Sharma PK, Sarma BK, Teli B, Bajpai R, Bhowmik A, Singh HV, Saxena AK. Inter-Genera Colonization of Ocimum tenuiflorum Endophytes in Tomato and Their Complementary Effects on Na+/K+ Balance, Oxidative Stress Regulation, and Root Architecture Under Elevated Soil Salinity. Front Microbiol. 2021; 12:744733. doi: 10.3389/fmicb.2021.744733.

37. Sahu PK, Singh S, Gupta AR, Gupta A, Singh UB, Manzar N, Bhowmik A, Singh HV, Saxena AK. Endophytic bacilli from medicinal-aromatic perennial Holy basil (Ocimum tenuiflorum L.) modulate plant growth promotion and induced systemic resistance against Rhizoctonia solani in rice (Oryza sativa L.). Biol Control. 2020; 150:104353. doi: 10.1016/j.biocontrol.2020.104353.

38. Adarsh A, Kumar R, Bhardwaj A, Chaudhary H. Correlation matrix study in bitter gourd for qualitative and quantitative traits. J Pharmacogn Phytochem. 2019; 8(3):3023–7.

39. Kołton A, Długosz-Grochowska O, Wojciechowska R, Czaja M. Biosynthesis Regulation of Folates and Phenols in Plants. Sci Hortic. 2022; 291:110561. doi: 10.1016/j.scienta.2021.110561.

40. Hamilton CE, Gundel PE, Helander M, Saikkonen K. Endophytic mediation of reactive oxygen species and antioxidant activity in plants: a review. Fungal Divers. 2012; 54:1–10.

41. Torres MS, White Jr JF, Zhang X, Hinton DM, Bacon CW. Endophyte-mediated adjustments in host morphology and physiology and effects on host fitness traits in grasses. Fungal Ecol. 2012; 5(3):322–30.

42. White Jr JF, Torres MS. Is plant endophyte-mediated defensive mutualism the result of oxidative stress protection? Physiol Plant. 2010; 138(4):440–6.

43. Hagaggi NSA, Mohamed AAA. Plant–bacterial endophyte secondary metabolite matching: a case study. Arch Microbiol. 2020; 202(10):2679–87. doi: 10.1007/s00203-020-01989-7.

44. Govindappa M, Channabasava R, Sowmya DV, Meenakshi J, Shreevidya MR, Lavanya A, Santoyo G, Sadananda T. Phytochemical Screening, Antimicrobial and in vitro Anti-inflammatory Activity of Endophytic Extracts from Loranthus sp. Pharmacogn J. 2011; 3(25):82–90. doi: 10.5530/pj.2011.25.15.

45. Banerjee S, Chanda A, Adhikari A, Das A, Biswas S. Evaluation of Phytochemical Screening and Anti Inflammatory Activity of Leaves and Stem of Mikania scandens (L.) Wild. Ann Med Health Sci Res. 2014;4(4):532–6. Epub 2014/09/16. doi: 10.4103/2141-9248.139302.

46. Mazumder K, Ruma YN, Akter R, Aktar A, Hossain MM, Shahina Z, Mazumdar S, Kerr PG. Identification of bioactive metabolites and evaluation of in vitro anti-inflammatory and in vivo antinociceptive and antiarthritic activities of endophyte fungi isolated from Elaeocarpus floribundus blume. J Ethnopharmaco. 2021; 273:113975. doi: 10.1016/j.jep.2021.113975.

47. Sharma G, Agarwal S, Verma K, Bhardwaj R, Mathur V. Therapeutic compounds from medicinal plant endophytes: molecular and metabolic adaptations. J Appl Microbiol. 2023; 134(4). doi: 10.1093/jambio/lxad074.

48. Hasan N, Farzand A, Heng Z, Khan IU, Moosa A, Zubair M, Na Y, Ying S, Canming T. Antagonistic potential of novel endophytic Bacillus strains and mediation of plant defense against Verticillium wilt in upland cotton. Plants. 2020; 9(11):1438.

49. Jasim B, Sreelakshmi K, Mathew J, Radhakrishnan E. Surfactin, iturin, and fengycin biosynthesis by endophytic Bacillus sp. from Bacopa monnieri. Microb Ecol. 2016; 72:106–19.

50. Abdelgalil SA, Kaddah MMY, Duab MEA, Abo-Zaid GA. A sustainable and effective bioprocessing approach for improvement of acid phosphatase production and rock phosphate solubilization by Bacillus haynesii strain ACP1. Sci Rep. 2022; 12(1):8926. doi: 10.1038/s41598-022-11448-6.

51. Masruroh IF, Sanjaya EH, Alvionita M, Suharti S, editors. Screening of Factors Influencing Keratinase Fermentation from Bacillus Haynesii BK1H using The Plackett-Burman Design (PBD). E3S Web Conf; 2024: EDP Sciences.

52. Bopaiah BBK, Kumar DAN, Balan K, Dehingia L, Reddy MKRV, Suresh AB, Nadumane VK. Purification, characterization, and antiproliferative activity of L-methioninase from a new isolate of Bacillus haynesii JUB2. J Appl Pharm Sci. 2020; 10(10):054–61.

53. Romanenko A. Enzymatic bioprospection of cultured bacterial diversity of Al-Wahbah crater, Saudi Arabia 2022. KAUST Research Repository. 10.25781/kaust-q8643.

54. Perez JL, Jayaprakasha G, Patil BS. Metabolite profiling and in vitro biological activities of two commercial bitter melon (Momordica charantia Linn.) cultivars. Food Chem (Oxf). 2019; 288:178–86. doi: 10.1016/j.foodchem.2019.02.120.

55. Wang X, Sun W, Cao J, Qu H, Bi X, Zhao YJ. Structures of new triterpenoids and cytotoxicity activities of the isolated major compounds from the fruit of Momordica charantia L. J Agric Food Chem. 2012;60(15):3927–33.

56. Muhammad Jihad S. Formulation of Momordica charantia fruit and Syzygium polyanthum leaf extracts based on in vitro antioxidant and inhibitory activity of α-amylase and α-glucosidase/Muhammad Jihad Sandikapura: University of Malaya; 2018.

57. Zhang R, Deng P, Dai A, Guo S, Wang Y, Wei P, et al. Design, synthesis, and biological activity of novel ferulic amide Ac5c derivatives. ACS omega. 2021;6(41):27561–7.

58. Liu H, Tang Y, Deng Z, Yang J, Gan D. Boosting the Antioxidant Potential of Polymeric Proanthocyanidins in Litchi (Litchi chinensis Sonn.) Pericarp via Biotransformation of Utilizing Lactobacillus Plantarum. Foods [Internet]. 2023; 12(12).

59. Zhu Y, Xie D-Y. Docking characterization and in vitro inhibitory activity of flavan-3-ols and dimeric proanthocyanidins against the main protease activity of SARS-Cov-2. Front Plant Sci. 2020; 11:1884.

60. Mamillapalli V, Shaik AR, Avula PR. Antiasthmatic activity of 2-piperidone by selective animal models. J Res Pharm. 2020; 24:334–40.

61. Ul Hassan Z, Al Thani R, Alnaimi H, Migheli Q, Jaoua S. Investigation and application of Bacillus licheniformis volatile compounds for the biological control of toxigenic Aspergillus and Penicillium spp. ACS omega. 2019; 4(17):17186–93.

62. Zhang Y, Cui J, Cao J, Pan H, Zhao Y. Novel Active Constituents of Momordica Charantia L. Plant Med. 2009; 75(04):S-43.

63. Chen B-S, Yang L-H, Ye J-L, Huang T, Ruan Y-P, Fu J, Huang PQ. Diastereoselective synthesis and bioactivity of long-chain anti-2-amino-3-alkanols. Eur J Med Chem. 2011; 46(11):5480–6. doi: 10.1016/j.ejmech.2011.09.010.

64. Zeece M. Chapter Seven - Food additives. In: Zeece M, editor. Introduction to the Chemistry of Food: Academic Press; 2020. p. 251-311.

65. Perez-Castillo Y, Lima TC, Ferreira AR, Silva CR, Campos RS, Neto JBA, Hemerson I. F. Magalhães HIF, Cavalcanti BC, Júnior HVN, de Sousa DP. Bioactivity and Molecular Docking Studies of Derivatives from Cinnamic and Benzoic Acids. Biomed Res Int. 2020; 2020:6345429. doi: 10.1155/2020/6345429.

66. Hoda S, Gupta L, Agarwal H, Raj G, Vermani M, Vijayaraghavan P. Inhibition of Aspergillus fumigatus Biofilm and Cytotoxicity Study of Natural Compound Cis-9-Hexadecenal. J Pure Appl Microbiol. 2019;13(2).

67. Rahman MM, Paul SI, Rahman A, Haque MS, Ador MAA, Foysal MJ, Islam MT, Rahman MM. Suppression of Streptococcosis and modulation of the gut bacteriome in Nile tilapia (Oreochromis niloticus) by the marine sediment bacteria Bacillus haynesii and Advenella mimigardefordensis. Microbiol Spectr. 2022;10(6):e02542–22.

68. Diomande SE, Nguyen-The C, Guinebretière M-H, Broussolle V, Brillard J. Role of fatty acids in Bacillus environmental adaptation. Front Microbiol. 2015;6:813.

69. Tanvir R, Sajid I, Hasnain S, Kulik A, Grond S. Rare actinomycetes Nocardia caishijiensis and Pseudonocardia carboxydivorans as endophytes, their bioactivity and metabolites evaluation. Microbiol Res. 2016;185:22–35. doi: 10.1016/j.micres.2016.01.003.

70. Dudek B, Rybka J, Bugla-Płoskońska G, Korzeniowska-Kowal A, Futoma-Kołoch B, Pawlak A, Gamian A. Biological functions of sialic acid as a component of bacterial endotoxin. Front Microbiol. 2022;13. doi: 10.3389/fmicb.2022.1028796.

